# *In vitro* and *in vivo* evaluation of the biofilm-degrading *Pseudomonas* phage Motto, as a candidate for phage therapy

**DOI:** 10.1101/2022.10.12.512010

**Authors:** Prasanth Manohar, Belinda Loh, Dann Turner, Ramasamy Tamizhselvi, Marimuthu Mathankumar, Namasivayam Elangovan, Ramesh Nachimuthu, Sebastian Leptihn

## Abstract

Infections caused by *Pseudomonas aeruginosa* are becoming increasingly difficult to treat due to the emergence of strains that have acquired multidrug resistance. Therefore, phage therapy has gained attention as an alternative to the treatment of pseudomonal infections. Phages are not only bactericidal but occasionally show activity against biofilm as well. Here, we describe the *Pseudomonas* phage Motto which has the ability to clear *P. aeruginosa* infections in an animal model and also exhibited biofilm-degrading properties. The phage has substantial antibiofilm activity against strong biofilm-producing isolates (n = 10), with at least a 2-fold reduction within 24 hours. To demonstrate the safety of using phage Motto, we performed cytotoxicity studies with human cell lines (HEK 293 and RAW 264.7 macrophages). Using a previously established *in vivo* model, we demonstrated the efficacy of Motto in *C. elegans*, with a 90% survival rate when treated with the phage at an MOI of 10.

**Importance:** Phages are often evaluated mainly on their ability to kill bacterial hosts. One important aspect, however, is often neglected: Their ability to degrade biofilms. Not all phages are able to disintegrate biofilms. While phages can kill planktonic cells, it also has often been observed that phages are not able to infect those that are embedded in biofilms. Phage Motto appears to have highly efficient enzymes that degrade biofilms, and might therefore be a highly valuable therapeutic candidate.

## Introduction

The nosocomial pathogen *Pseudomonas aeruginosa* belongs to the ESKAPE group of bacteria which poses a serious threat to public health. As the rates of antibiotic resistance increase, *P. aeruginosa* infections are becoming more difficult and sometimes impossible to treat, especially due to their tendency to form biofilms in chronic infections. Phage therapy, the use of bacterial viruses to kill bacterial pathogens, might offer a solution (1-4).

Lytic bacteriophages (phages) are bacterial viruses that replicate inside a suitable host and get released through lysis which destroys the bacterial cell (5). Phage therapy case studies and clinical trials have been increasing over the last decade as they are often seen as the only suitable alternative to conventional antibiotic therapy, commonly used additionally due to synergetic effects (6-8). However, clinical trials have produced mixed results due to the specificity of phages and the rapid development of phage resistance (9).

As one of the ESKAPE pathogens (which include *Enterococcus, Staphylococcus, Klebsiella, Acinetobacter, Pseudomonas* and *Enterobacter*), *P. aeruginosa* is a common pathogen known to cause hospital- and community-acquired infections and is a notorious complication in cystic fibrosis patients (10,11). The increasing rate of antibiotic-resistant *P. aeruginosa* strains is a major concern, as they are attributed to high mortality rates in healthcare-associated infections (12-14). Moreover, the ability to form extensive biofilms is a major virulence determinant during pathogenesis, especially in chronic infections (15). Due to their complex composition, biofilms are able to prevent efficient diffusion of antibiotics to the target cells, making treatment even more challenging even if the cells are susceptible when not embedded in this polymeric multicomponent matrix. Such biofilm-forming pathogenic strains are often found in cystic fibrosis patients, but can also cause ventilator-associated pneumonia (16). However, available anti-pseudomonal antibiotics have ceased to be effective in an increasing number of cases as many strains have acquired antibiotic-resistant genes (17). Therefore, an alternative therapy is needed against pseudomonal infections that also show activity against biofilm-forming cells. This is particularly important, as while some phages can be highly efficient in killing planktonic cells, biofilms can act as a protective layer to prevent phage infection of embedded bacteria (18,19).

In this study, we describe the isolation and characterisation of a novel *Pseudomonas* phage, Motto. Motto has a broad host range infecting more than 50% of the clinical strains tested and can efficiently disrupt biofilms formed by the pathogen. Furthermore, safe and efficient use was demonstrated by cytotoxicity assays in human cell cultures and in the animal model *C. elegans*.

## Materials and methods

### Bacterial strains

The bacterial strains used in this study were collected from the Hi-tech diagnostic centre, in Chennai. A total of 50 distinct *P. aeruginosa* isolates collected from clinical samples were used in this study. All the isolates were previously characterized and reported (20,21). The isolates were tested for susceptibility to cefotaxime, ciprofloxacin, gentamicin, meropenem and tetracycline using the micro-broth dilution-MIC method following Clinical Laboratory Standard Institute (CLSI) guidelines (22). The antibiotic sensitivity results were recorded in accordance with CLSI guidelines.

### Sample collection and isolation of bacteriophages

For the isolation of bacteriophages, sewage water samples were collected from different parts of Tamil Nadu, in Chennai (Cooum River, 13.0827° N, 80.2707° E), Vellore (municipal sewage water system and hospital sewage water, 12.9165° N, 79.1325° E), and Karur (sewage treatment plant, 10.9601° N, 78.0766° E) and Coimbatore (domestic sewage wastewater, 11.0168° N, 76.9558° E). The sewage water samples were collected in containers (up to 1 L), transported to the laboratory and stored at 4°C. For the isolation of bacteriophages, the phage enrichment method was followed as described previously (23). Briefly, to the 10 mL of exponentially grown bacterial (host) culture, 30 mL of sewage water sample was added and incubated at 37°C for 20 hours. Then, the mixture was centrifuged at 6,000 × *g* for 15 min and the supernatant was collected. The collected phage lysate was filtered through 0.22-micron syringe filters (≈ 2 mL) and tested for the presence of bacteriophage using spot test and double agar overlay methods (23).

Briefly, in the spot test, a bacterial lawn was prepared using the host bacteria on the Luria Bertani (LB) agar and 10 µL of phage filtrate was spotted. The plates were incubated at 37°C for 16 hours. The clearance of bacterial growth on the spotted areas denotes the presence of phage activity. For the double agar overlay method, bacteria (200 µL) and phage filtrate (100 µL) were mixed and kept undisturbed for 15 min. Then, soft agar (0.75% w/v, 3 mL) was added, mixed and poured onto pre-prepared LB agar plates. The plates were incubated at 37°C for 16 hours and the appearance of plaques represents the phage lytic activity. For the screening of bacteriophages, a single plaque was chosen and purified, and the double agar overlay method was repeated thrice.

### Precipitation and purification of bacteriophages

The bacteriophages were precipitated using the polyethylene glycol-sodium chloride (PEG-NaCl) method (23). Briefly, 1 M PEG and 0.1 M NaCl were added to the phage filtrate, mixed gently and incubated at 4°C for 24 hours. The precipitant was centrifuged at 15,000 × *g* for 30 min and 100 µL SM buffer [For 1 L: 5.8 g of NaCl; 50 mL of 1 M Tris-HCl [pH 7.5]; 2 g of MgSO_4_.7H_2_O] was added to the pellet. The phage precipitant was stored at −20°C. To purify the bacteriophages, the sucrose gradient method was used. In brief, varying concentrations of sucrose solution were prepared from 12.5 to 52.5% w/v and layered in descending order of concentration. One mL of phage suspension was added at the top and centrifuged at 30,000 × *g* for 1 hour resulting in the phages being concentrated as a visible band. The phage band was removed carefully using a micropipette, dialyzed and the purified phages were stored at −20°C.

### Host range analysis

To study the infectivity of the isolated *Pseudomonas* phage against different bacteria, the selected phage was tested against other 49 strains of *P. aeruginosa* (plus one serving as host), five strains of *E. coli*, three strains of *K. pneumoniae* and three strains of *A. baumannii*. Briefly, the spot test was performed using the phages at 10^3^ PFU/mL against the test bacteria. The lysis of bacteria and the appearance of clear zones represent phage activity. The isolates with the positive spot test results were further chosen for the double agar overlay method. The appearance of plaques indicated phage lytic activity.

### Life cycle studies

The phage life cycle studies include three different stages; adsorption, latent period and burst size. To determine the adsorption time, the bacterial cells at 10^6^ CFU/mL were mixed with the phages at the multiplicity of infection of 0.01 (MOI) and incubated at 37°C. From the mixture, 100 µL was removed at every 5 min interval for 45 min and diluted in 4.9 mL of LB broth. After incubating for 30 min at 37°C, the non-adsorbed phages were determined using the double agar overlay method. The adsorption curve was plotted based on the number of non-adsorbed phages against time.

A one-step growth experiment was used to study the latent period, the time taken for the phages to multiply inside the bacteria, and burst size, the number of phages released from each infected cell. Briefly, the bacterial cells (10^6^ CFU/mL) were mixed with the phages at an MOI of 0.01 and the phages were allowed to adsorb for 30 minutes at 37°C. Then, the mixture was centrifuged at 12,000 × *g* for 5 min and to the pellet; 10 mL of LB broth was added and incubated at 37°C. Then, at every 10 min interval, the samples were taken and titrated against the host bacterium. Both the latency period and burst size were plotted against time (in min). All the data are presented as mean ± standard deviation (SD) of at least three independent experiments.

### Stability studies

The stability of the phages was determined at different pH (1-12) and temperatures (20-70°C), following previously described protocols (24). Briefly, the phage suspension adjusted to a titer of 10^6^ PFU/mL was incubated under various physical conditions. The reduction in phage titer was analysed after 60 minutes by overlay plaque assay. In brief, the pH stability studies were performed using SM buffer and the phage lysates were incubated at varying pHs ranging from 1 to 12. For thermal stability studies, the phage lysates were incubated at 20 to 70°C for 60 min and tested for phage activity. All the data are presented as mean ± standard deviation (SD) of at least two independent experiments.

### Transmission electron microscopic (TEM) analysis

To morphologically characterize the bacteriophage structure, the phages were negatively stained using uranyl acetate and visualized under TEM (FEI-TECNAI G2-20 TWIN, Bionand, Spain) at the VIT-TEM facility (23). Briefly, in a copper grid, 2 µL of purified phages at 10^8^ PFU/mL was added and allowed to adsorb for 10 min. The excess samples were removed and the grid was dried. Staining was performed by the addition of 2% (w/v) uranyl acetate. The copper grid was washed three times with distilled water to remove excess stain and allowed to dry for 30 min prior to examination.

### Whole Genome Sequencing and analysis

Initially, the phage DNA was isolated using the phenol-chloroform method (24:1) and precipitated using ethanol (25). The Nextera XT DNA library preparation kit was used to create the sequencing libraries. The phage genome was then sequenced using the Illumina Hiseq platform. A total of 5,885,529 clean reads of 150 bp (paired-end format) were used to assemble the genome. The short-read sequence data were assembled using Unicycler (v0.4.7) (26). The assembly was performed after quality filtering; quality control employing FastQC, MultiQC, and trimmomatic (27-29) and completion of the assembled genome was determined, and the coverage and depth were calculated by bedtools (30). Genome annotation was completed using Prokka 1.14.5 and Galaxy-Apollo (31,32). All tools were run with default parameters unless otherwise specified.

### Comparative genomics and phylogenetics

All phages classified as species within the family *Drexlerviridae* in the ICTV virus Metadata Resource version 37 (https://ictv.global/vmr) were retrieved from GenBank. Nucleotide sequence similarity between phages was compared using VIRIDIC with default settings. To determine appropriate core genes conserved within the family for phylogenetic analysis, phage genomes were first reannotated with Prokka version 1.14.5 (31) with a custom hmm pVOGs database to provide consistent gene calls. Pangenome analysis was performed with the predicted protein-coding sequences using PIRATE (33) with parameters of 35, 45 and 55% identity. Maximum likelihood phylogenetic trees were constructed from ClustalO (34) alignments with IQTree2 (35). IQTree2 was run with ModelFinder (36) to select the most appropriate evolutionary model according to the Bayesian information criterion with 1000 Ultra-fast bootstrap replicates (37) and the SH-aLRT test. Trees were rooted using an outgroup (*Dhillonvirus* JG1), before being annotated with ITOL (38).

### Biofilm clearance assay

To study biofilm formation and the anti-biofilm activity of phages, a microtiter plate assay was performed. Briefly, the *P. aeruginosa* strains (n=32, based on phage activity) were grown in a microtiter plate in LB broth for 24 hours. The samples were then stained using crystal violet (CV) and washed, before the OD at 595 nm was read (BioTek, India). Biofilm formation was recorded as weak, moderate and strong based on the OD values in comparison to the control. (Range: weak with OD of < 1.0, moderate with OD of 1.0-2.0, strong with OD of > 2.0).

To study the anti-biofilm activity of phages, 10 strong biofilm-forming (based on OD) *P. aeruginosa* isolates were chosen (*P. aeruginosa* strains 01, 08, 11, 16, 27, 32, 35, 37, 42, 47). Anti-biofilm activity of the phages was determined after adding the phages (10^6^ PFU/mL) to the biofilm. After 24 hours of incubation at 37°C, the CV-staining protocol was performed and the OD values were determined. All the data are presented as mean ± standard deviation (SD) of at least three independent experiments.

### Cytotoxicity of bacteriophage on mammalian cells: Maintenance of mammalian cell lines

The human embryonic kidney (HEK 293) cell lines were obtained from the VIT Cell Culture facilities, SBST (39). The cell lines were grown in Dulbecco’s modified eagle’s medium (DMEM) containing 10% (v/v) fetal bovine serum (FBS) and 5 mL/L (100x) antibiotic solution, which contains 100 U/mL penicillin, and 100 mg/mL streptomycin, and 25 μg/mL Amphotericin B. Cells were maintained at 37°C in 5% atmospheric CO_2_ and 95% air. The cells were grown to 80-90% confluence in a T25 flask. Another cell line, RAW 264.7 macrophage cells (established murine macrophage cell lines, ATCC) were cultured in complete DMEM with high glucose and supplemented with 10% (v/v) FBS, 100 U/mL penicillin, and 100 mg/mL streptomycin, pH 7.4 at 37°C in 5% CO_2_ until 70 to 80% confluence. Cells (3’10^5^ cells/mL) were then cultured overnight for further assays.

### Testing the bacteriophage-mammalian cell line interaction

The purified bacteriophages at 10^6^ PFU/mL were used to treat the mammalian cell lines, HEK 293 and RAW 264.7 macrophages to determine the cytotoxicity (40). Briefly, 5×10^5^ cells were seeded into wells of a six-well plate and maintained at 37°C, 5% in DMEM with 10% FBS. After 24 hours, cells were washed with PBS and treated with phage suspension without FBS. HEK293 cells and RAW 264.7 macrophages treated with Triton X-100 served as a positive control. To count the cells using a haemocytometer, 100 μL of cell suspension was taken and 100 μL of 0.4% trypan blue was added. A 20 μL of cell suspension was filled under the cover glass and the cells were counted under 10x magnification.

### Testing the efficacy of phage in the animal infection model C. elegans

#### C. elegans maintenance

Bristol N2 (wild-type) *C. elegans* was used in this study, and the nematode was maintained and propagated on nematode growth media (NGM; 17 g agar, 3 g NaCl, 2.5 g peptone, 0.5 mL of 1 M CaCl_2_, 1 mL of 5 mg/mL cholesterol, 1 mL of 1 M MgSO_4_, 25 mL KH_2_PO_4_ buffer [pH 6.0] per litre) plates which carried *E. coli* OP50 as a source of food at 20°C by standard protocols (41). Age-synchronised L4 worms were exposed to TSB, *P. aeruginosa* and *Pseudomonas* phage in a 96-well microtiter plate containing M9 buffer in each of the wells for the phage efficacy assay.

### Phage efficacy assay

A liquid-based assay to study the efficacy of phages against pathogenic bacteria was previously developed by our research team (42). To test the efficacy of *Pseudomonas* bacteriophage in recovering the nematodes from pseudomonal infection, two types of studies were performed i.e., therapeutic treatment and prophylactic treatment. For a liquid-based assay, a 96-well microtiter plate was filled with M9 buffer to which overnight culture of *E. coli* OP50 or an equivalent amount of *P. aeruginosa*, with or without phage was added. Then 20 mature L4 nematodes were transferred into the solution (42). The total volume in the microtiter plate was maintained at 100 μL. To test the phage efficacy, different concentrations of *P. aeruginosa* were used to test the infectivity. Accordingly, overnight cultures of bacteria were diluted to 10^5^ CFU/mL (based on our previous study), and the efficacy of phage treatment was studied using varying concentrations of phages, i.e., 10^5^, 10^6^ and 10^7^ CFU/mL. Therefore, the study includes bacteria and phage concentrations at 1:1 (10^5^ CFU/mL: 10^5^ PFU/mL), 1:10 (10^5^ CFU/mL: 10^6^ PFU/mL) and 1:100 (10^5^ CFU/mL: 10^7^ PFU/mL).

Group 1 (control) consisted of M9 buffer (60%) with *E. coli* OP50 at 10^5^ CFU/mL (40%) and 20 nematodes. Group-2: M9 buffer, 20 nematodes and no bacteria. Group 3: M9 buffer (60%), TSB (40%) and 20 nematodes. Groups 2 and 3 were experimental controls. Group 4 (infection control): M9 buffer (60%), *P. aeruginosa* (40%) and 20 nematodes. Group 5 (heat-inactivated bacteria): M9 buffer (60%), heat-inactivated P. aeruginosa, 40% and 20 nematodes. Group 6 (phage toxicity test): M9 buffer (60%), *Pseudomonas* phage Motto (40%) and 20 nematodes. Group 7 (therapeutic treatment group): M9 buffer (40%), *P. aeruginosa* (30%), 20 nematodes; *Pseudomonas* phage Motto (30%) was added 2 h of exposure to bacteria. Group 8 (prophylactic treatment group): M9 buffer (40%), *Pseudomonas* phage Motto (30%), 20 nematodes; *P. aeruginosa* (30%) were added after 1 h. The plates were incubated at 20°C, and the survival of the nematodes was monitored every 24 hours for 5 days. The results were evaluated based on live nematodes (moving) and dead nematodes (lack of movement). All the experiments were repeated three times for statistical significance.

To analyze the presence of phages inside the nematodes or uptake of phages by the nematodes (42), phage numbers were determined as follows: Briefly, after 4 days, 10 nematodes were removed from groups 5, 6 and 7, which are phage only, therapeutic treatment and prophylactic treatment, respectively, to determine the phage titer using the double agar overlay method. In brief, the nematodes were washed thrice with buffer, vortexed, ground (with mortar and pestle), and centrifuged at 10,000 ’ *g* for 5 min. The supernatant was used to determine the phage titer (42).

### Statistical analysis

All the experiments were performed at least three times for statistical significance and the data are presented as the mean ± standard deviation (SD). For nematode studies, survival curves were plotted using the Kaplan-Meier method, and the log-rank test was used to calculate the difference in survival rates using GraphPad Prism software 9.0 (GraphPad Software, Inc., La Jolla, USA). *P < 0*.*05* was considered statistically significant using the log-rank test.

## Results

### Isolation of Pseudomonas infecting phages

For this study, we first screened for phages by using 50 clinical *P. aeruginosa* isolates with a mucoid phenotype as “bait”; most of the isolates were found to be multi-drug resistant (S.table 1). From five different environmental water samples, three phages infecting three different hosts were isolated and chosen for further studies. When analysing the host range of the phages, we found that one had the ability to infect the majority of tested strains 32/50 (64%) (S.fig.1). This phage was named Motto and was isolated from Cooum river water in Chennai, India. The plaque morphology was identical in all strains, forming clear plaques without any halo of 1.5-2 mm after overnight incubation (Fig.1). In our case, the exposure to phage Motto resulted in complete lysis of host bacteria on solid and in liquid media for all phage-sensitive isolates (n = 32); on plates, no growth was observed and in liquid, visibly clear cultures were seen, even after extended incubation times. Under the conditions we tested, we did not observe phage-resistant phenotypes.

### Pseudomonas phage Motto morphology is T1-like

Negative staining and TEM analysis allow the morphological characterization of phages. Using this technique, we found Motto to possess a siphovirus morphology with a long, flexible tail similar, that is morphologically similar to the members of the family *Drexlerviridae*. The icosahedral head is symmetrical and about 57 ± 1 nm in size, while the long non-contractile tail is 255 ± 1.5 nm in length (Fig.1).

**Figure 1:**
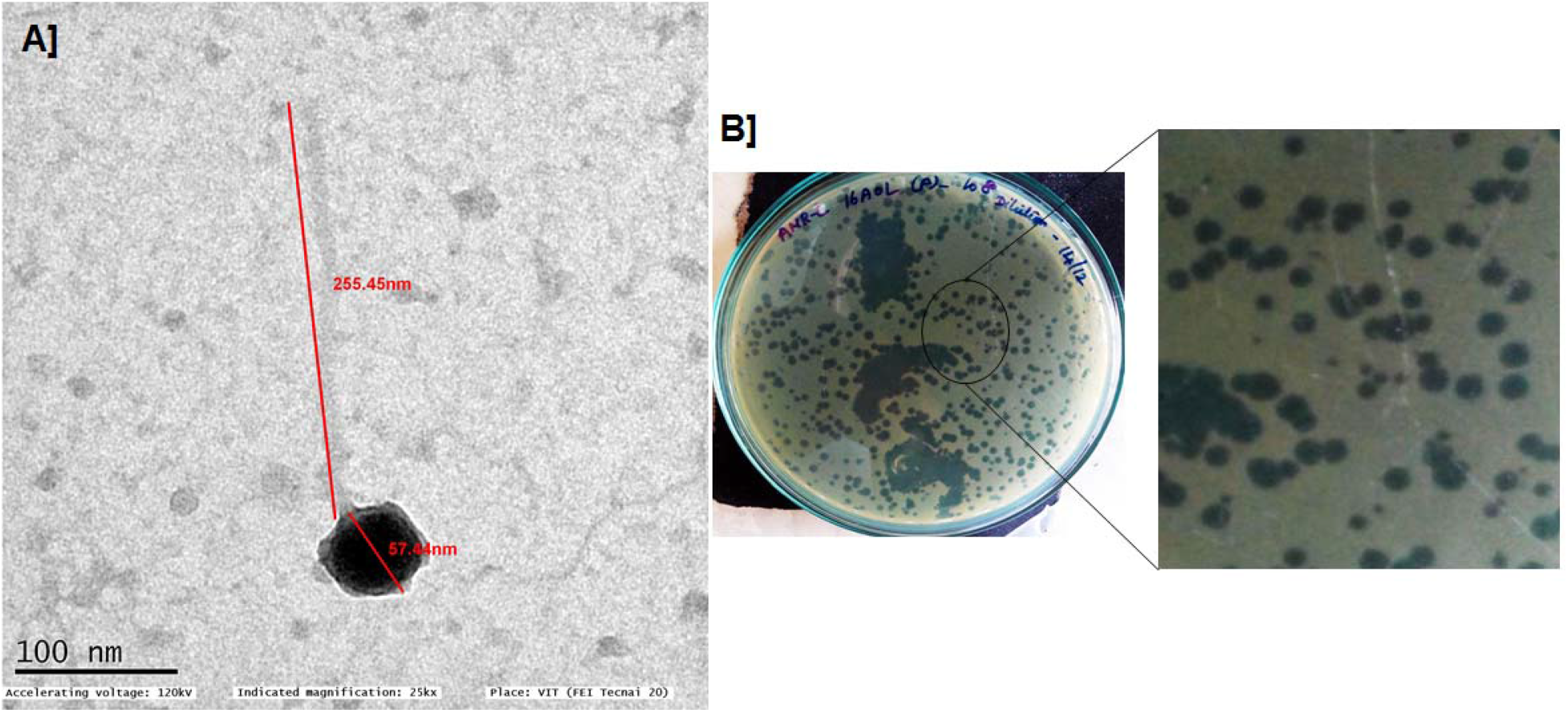
Transmission electron micrograph of *Pseudomonas* phage Motto (A) and morphology of plaques formed against *P. aeruginosa* isolate PA01 (B). TEM morphology shows that the phage exhibits a siphovirus morphology with an icosahedral head and long non-contractile tail.

### Replication and adsorption of phage Motto are rapid processes

The adsorption kinetics of *Pseudomonas* phage Motto to the host surface was rapid, with around 50% of the phages attaching to the host surface within approximately 10 minutes and 90% of phages being adsorbed to host cells within ∼25 min (Fig.2A). After adsorption, phage and host machinery facilitate the translocation of the viral genome followed by the replication of the virus. In the case of phage Motto this process requires approximately 30 minutes at 37°C (Fig.2B). Complete lysis of the culture, producing a transparent lysate, was observed at around 40 minutes. The burst size was calculated with 80 phage particles per host cell on average.

**Figure 2:**
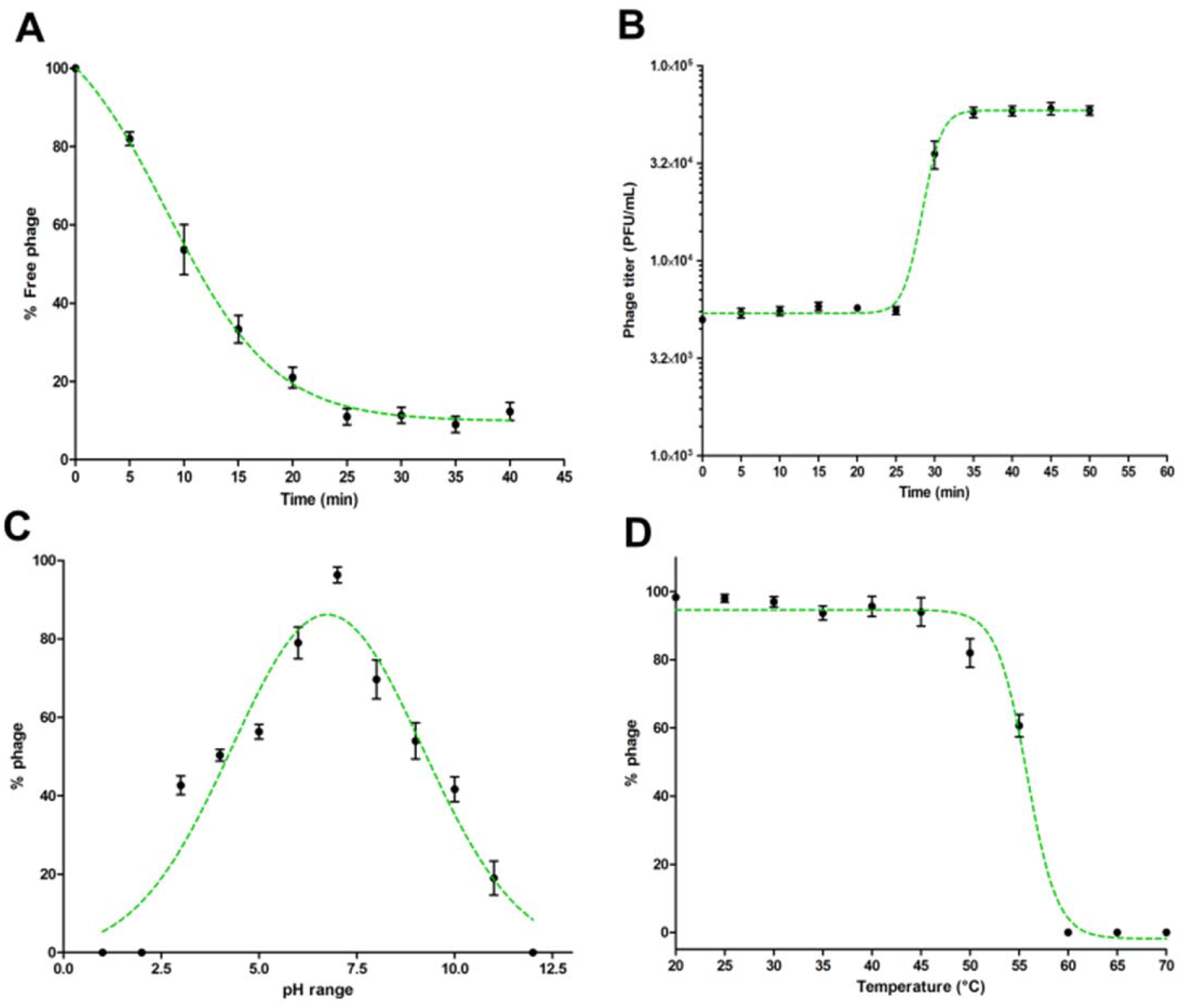
Characterization of *Pseudomonas* phage Motto. (A) Adsorption of phage to its host *P. aeruginosa*. The graph represents the % reduction in phage titer against the time (in min). The adsorption plots were fitted using an exponential decay reaction. (B) One-step growth curves of *Pseudomonas* phage Motto. The graph represents the change in titer against the time (in min). The plots were fitted using the sigmoidal fit. (C) pH stability of *Pseudomonas* phage Motto; the graph represents the % stability at varying pH conditions. The optimal pH stability was determined by plotting in the Gaussian fitting function. (D) Thermal stability of *Pseudomonas* phage Motto; the graph represents the stability at different temperatures (°C). The stability plots were fitted using the sigmoidal fit.

### Motto is stable under non-physiological conditions

Therapeutic phages have to be stable for storage and application. Therefore, we studied Motto’s stability at elevated temperatures and non-neutral pH values. We found phage Motto to be active at pH ranges between 3 and 11 with optimum stability at pH 6.7 (Fig.2C). The thermal stability showed that Motto is stable up to 55°C when incubated for 1 hour but completely inactivated at 60°C (Fig.2D).

### Genomic characterization of phage Motto

The Motto genome is a linear double-stranded DNA molecule with a size of 49,960 base pairs and a G+C content of 45% (NCBI accession number is ON843697) (Fig.3). Blastn was used on the entire genome sequence and the two closest relatives of Motto in the NCBI database are *Vibrio* virus 2019VC1 (NC_054898.1, Query coverage: 43%, sequence identity 88%) and *Salmonella* virus STSR3 (MT500539.1, Query coverage: 65%, sequence identity 82%) (Fig.3). We identified 84 predicted open reading frames (ORFs) and no tRNAs using Aragorn (43). More than half of the ORFs could not be assigned to any known protein, with 46 “hypothetical proteins” annotated whose functions remain to be investigated. S.table 2 shows the genes coding for proteins with putative functions, separated by their role. No depolymerase genes or biofilm-degrading hydrolases could be identified in Motto’s genome. No lysogeny-related genes were identified, confirming that Motto is lytic, while we also did not find any virulence factors in the genome.

**Figure 3:**
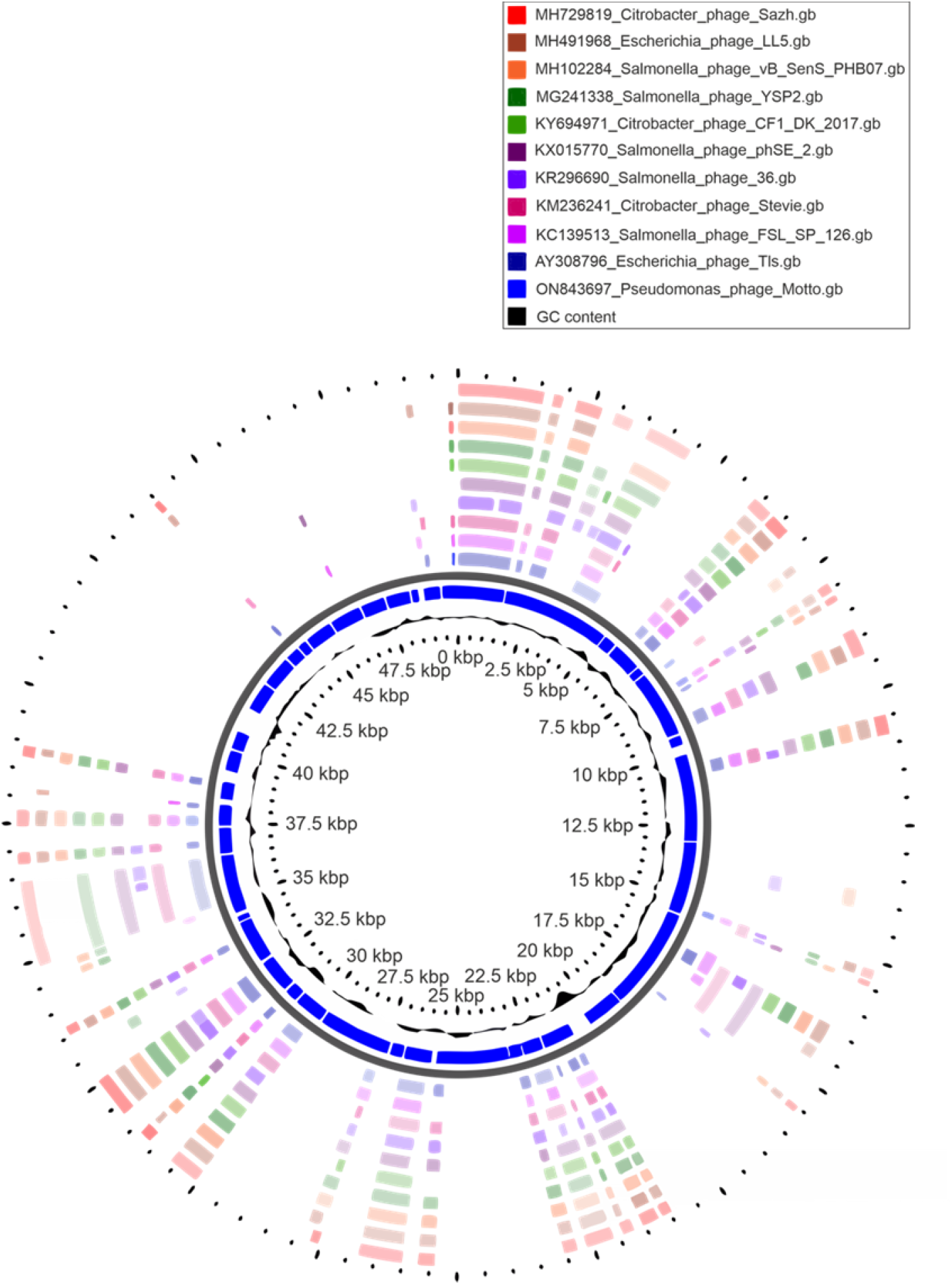
Circular genome representation of *Pseudomonas* phage Motto with the ten closest related phages using CGViewer.

### Taxonomic classification of Pseudomonas phage Motto

Initial analysis with BLASTn identified similarities to members of the *Tlsvirus*, family *Drexlerviridae*. To assess this relationship nucleotide sequences of phages classified within the family were subjected to pairwise comparison using VIRIDIC (S.Fig.2). Analysis of the nucleotide similarity between Motto and 167 species assigned to this viral family recapitulates the assigned genera and indicates that Motto represents a new species and genus according to current ICTV demarcation criteria (44).

Pangenome analysis revealed that a total of 10 gene products are conserved in all members of the *Drexlerviridae*. This number is likely to be greater when taking into account that the sequences are not colinear and that some genes were not called due to being split across the genome ends. Of the 12,363 proteins, 12,166 formed a total of 311 groups consisting of two or more representative sequences and 197 were unique singletons. Motto shares a maximum of 75% of proteins with *Klebsiella* phage GML-KpCol1 (MG552615) and a minimum of 44% with other *Drexlerviridae* species.

For phylogenetic analysis (Fig.4), a partitioned maximum-likelihood tree was constructed using 10 gene products conserved in all *Drexlerviridae* members; the small and large terminase subunits, major capsid protein, major tail protein, tape-measure, tail assembly protein, tail completion protein, tail tip protein and tail terminator protein. The topology of the tree revealed clades that were largely congruent with defined genera and subfamilies. Some exceptions were noted; firstly, the genus *Kyungwonvirus* comprising *Cronobacter* phages Esp2949-1 (JF912400) and CS01 (MH845412), separates the *Tempevirinae*. Secondly, the tree topology does not appear to support the inclusion of *Shigella* phage Sd1 (MF158042, *Wilsonroadvirus*) within the *Rogunavirinae* subfamily. The taxonomy of this family will require reassessment in the near future, to include new isolates deposited in the INSDC since its inception in 2019 (ICTV proposal 2019.100B; https://ictv.global/ictv/proposals/2019.100B.zip) and to allow a more detailed examination of the minor discrepancies observed here. Collectively, based on nucleotide similarity, a number of shared proteins and phylogenetic analysis, we propose that Motto represents a new species and genus within *Drexlerviridae*.

**Figure 4:**
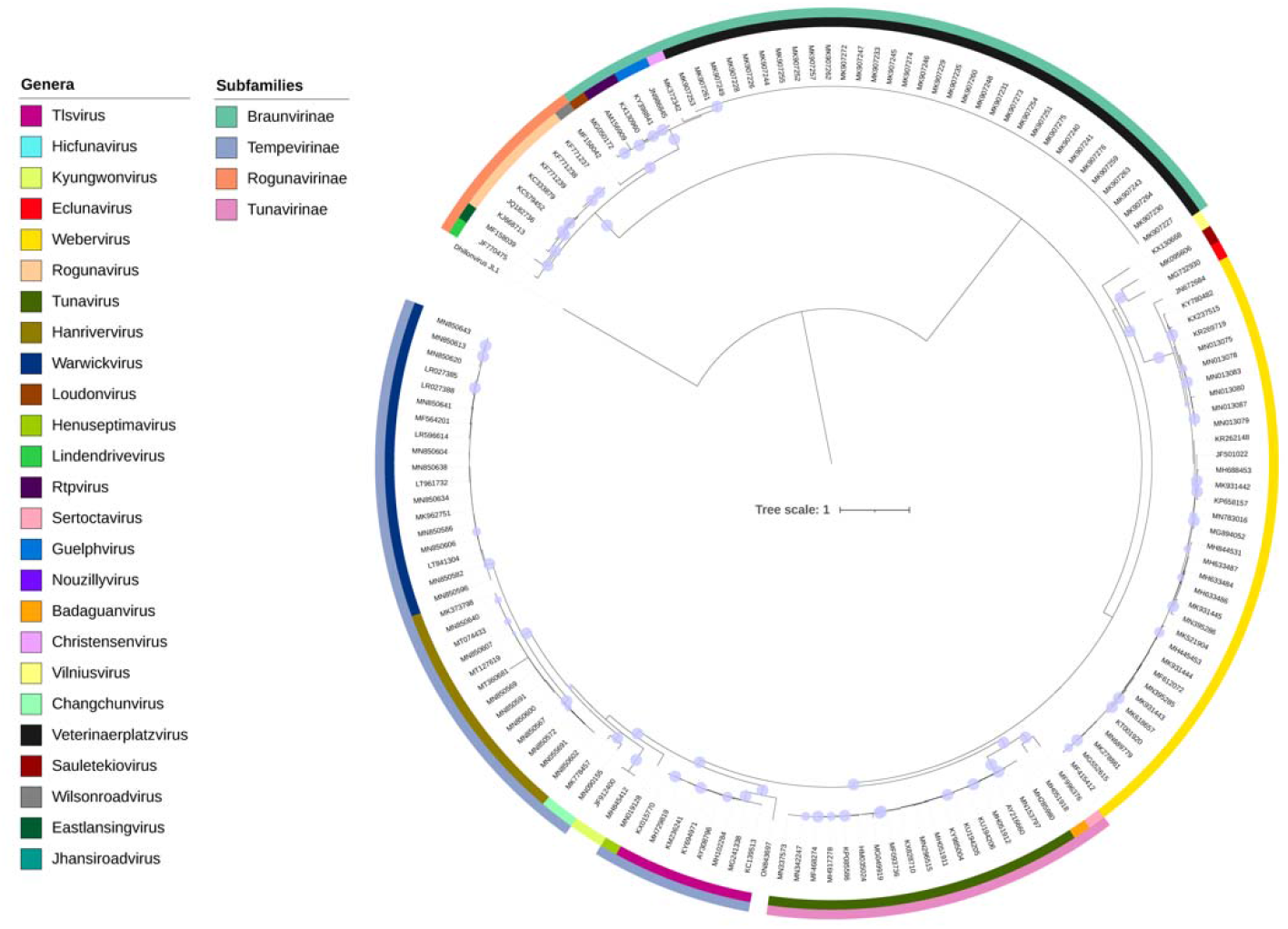
Maximum likelihood phylogenetic tree of proteins conserved across *Drexlerviridae*. Models were predicted for each alignment (TerS, Q.yeast+I+G4; TerL, LG+G4; major capsid protein, WAG+R3; tail assembly, LG+G4; tail completion protein, WAG+R4; tail tip protein Q.pfam+F+R5; major tail protein, WAG+G4; tail terminator protein, WAG+G4; helicase, WAG+G4 and tape measure protein, Q.pfam+F+R4). Ultrafast bootstrap support (1,000 replicates) >=95% are shown as filled circles. Colours on the inner and outer rings denote genera and subfamilies according to the key.

### Phage-mediated reduction of biofilms

The *in vitro* biofilm assay using the microtiter plate method showed that 10/32 isolates were strong biofilm producers (*P. aeruginosa* strains 01, 08, 11, 16, 27, 32, 35, 37, 42, 47). Of the 10 *P. aeruginosa* isolates tested, biofilm formation was substantially reduced in all and at least 75% reduction was observed (Fig.5). Importantly, our study showed the reduction of strong biofilms after 24 hours of treatment.

**Figure 5:**
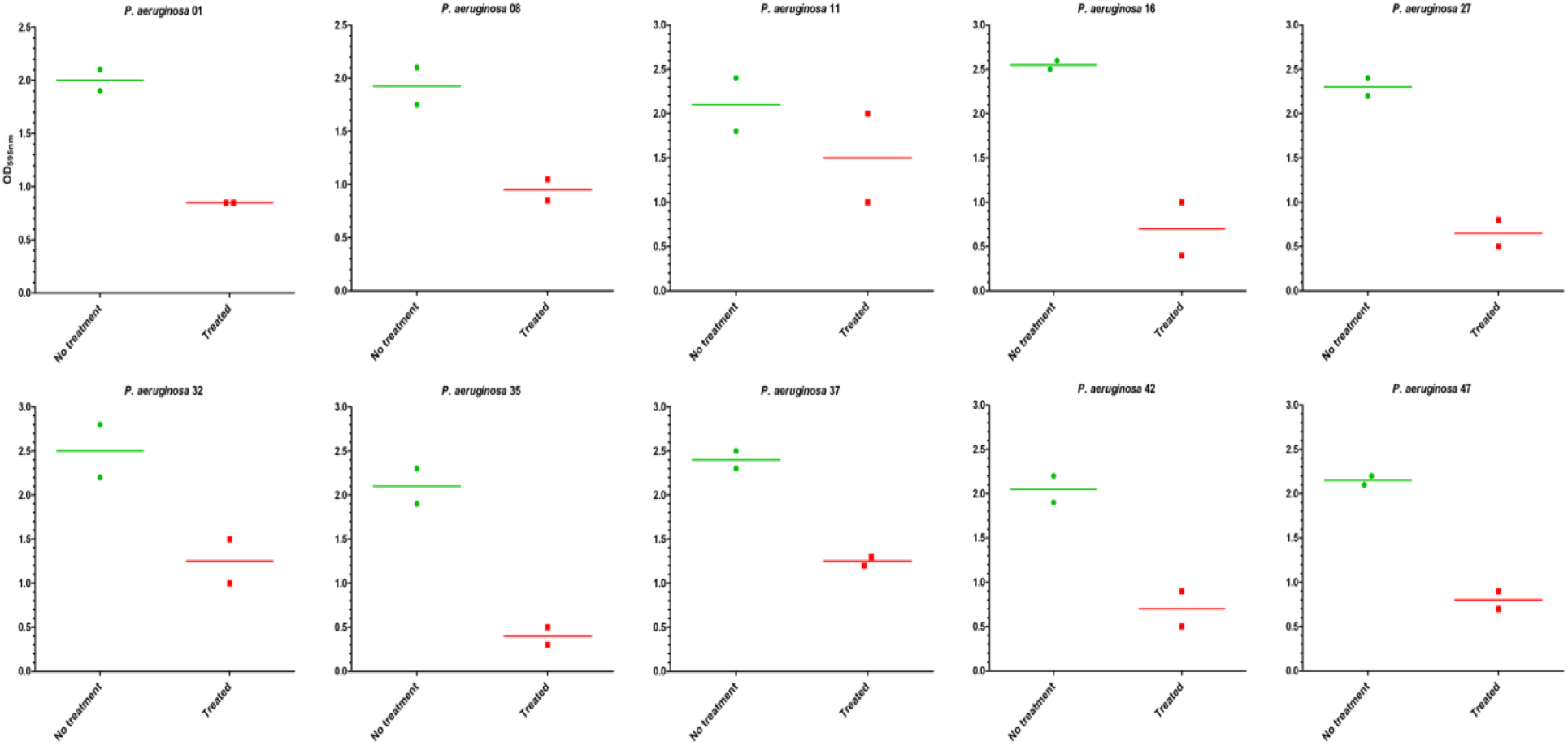
Anti-biofilm activity of *Pseudomonas* phage Motto against different strains of *P. aeruginosa*. A total of 10 strong biofilm producers were chosen and treated with phage (10^6^ PFU/mL) for 24 hours. The represented results are the OD_595nm_ values measured using a microtiter plate reader. The results were compared between the untreated and treated groups.

### Motto does not exhibit cytotoxic effects when applied to mammalian cells

*Pseudomonas* phage Motto was tested for toxicity with the mammalian cell lines, HEK 293 and RAW 264.7 macrophages. When the cells were exposed to a solution containing phages, no cytotoxic effect was observed using cell viability counts (Fig.6A, 6B). Phage-treated cells and normal cells exhibited no difference in cell morphology (Fig.6C,6D) after 6 and 24 hours of incubation.

**Figure 6:**
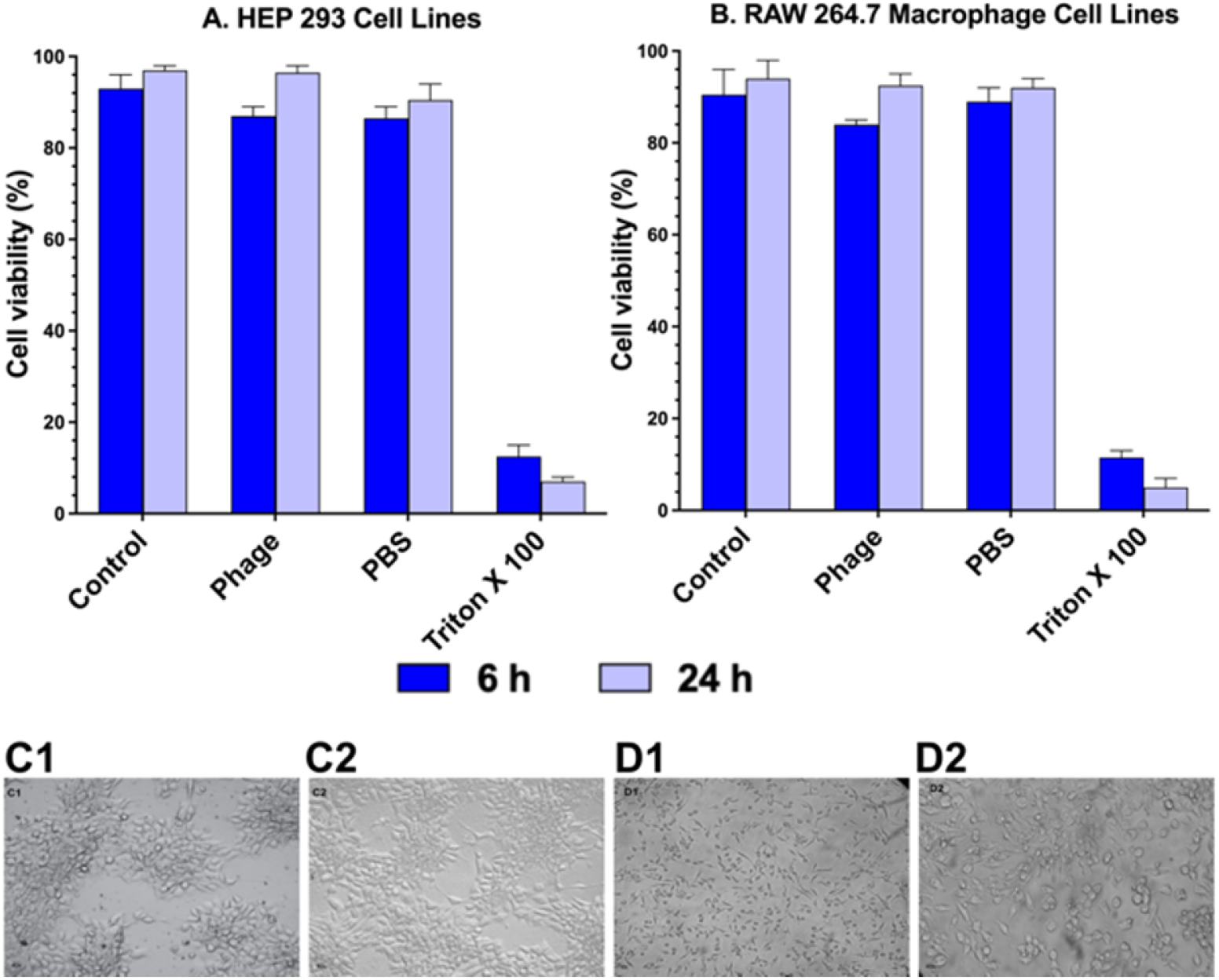
Cytotoxicity of *Pseudomonas* phage Motto on mammalian cells. The cytotoxicity was studied using HEP 293 and RAW 264.7 macrophage cell lines (5×10^5^ cells/well) using cell viability studies. The graphical representation of the viability of mammalian cells with and without phage treatment shows that the phage treatment did not affect the viability in comparison to the positive control (1% Triton X-100). (C - HEK 293 cell images, C1- Control, C2- Phage treated and D - RAW 264.7 cell images, D1- Control, D2- Phage treated). Cell images show there is no change in morphology after phage treatment (C2 and D2) as compared to controls.

### Application of Motto to nematodes results in reduced mortality

To evaluate the efficacy of *Pseudomonas* phage Motto against pathogenic pseudomonal infections, we employed the recently established *C. elegans* liquid-based assay (42). In the infection control group, all the nematodes died within 4 days (Fig.7A).

**Figure 7:**
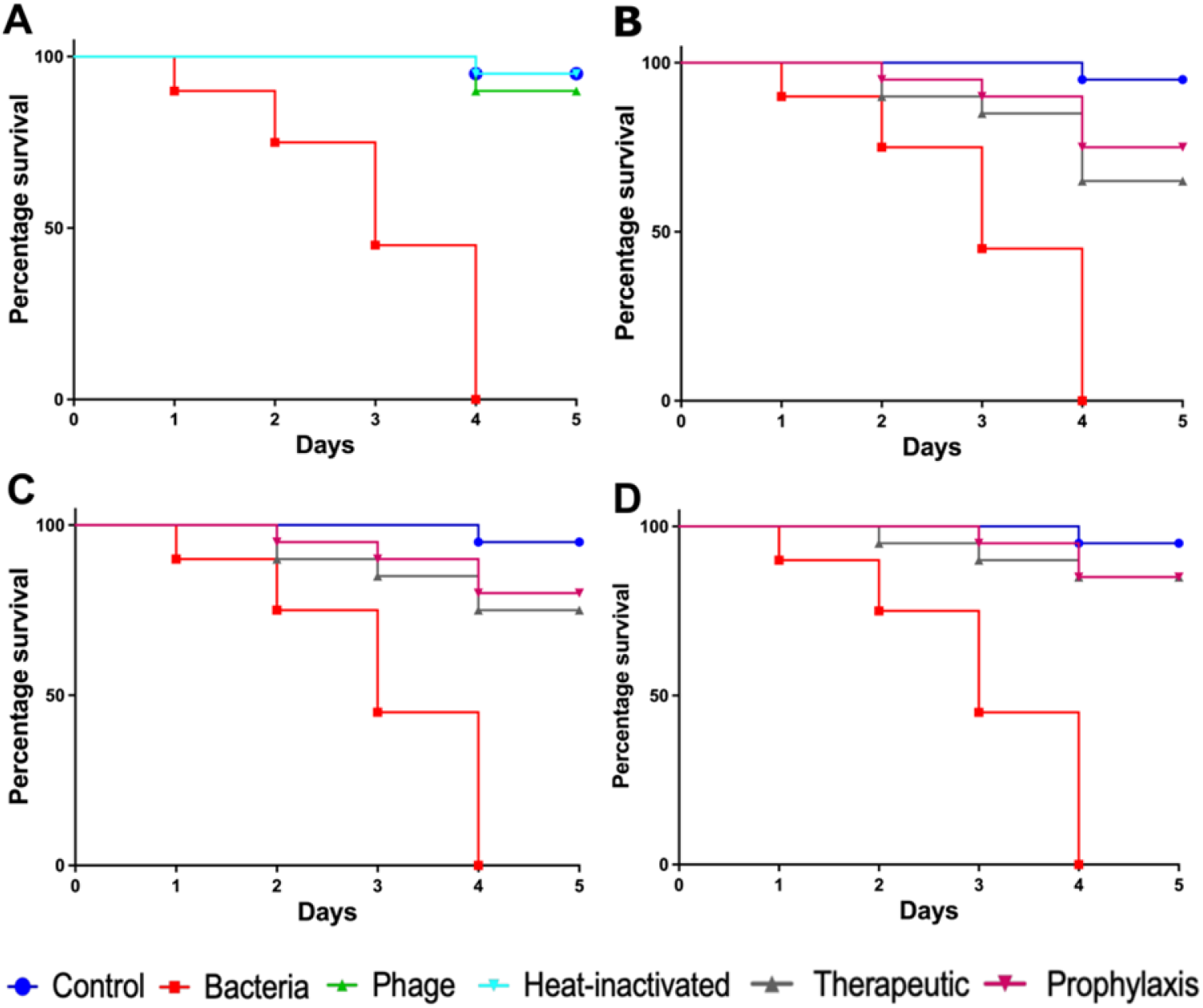
Pathogenicity of *P. aeruginosa* in *C. elegans* and efficacy of *Pseudomonas* phage Motto against pseudomonal infections. The control group consisted of *C. elegans* fed with *E. coli* OP50 and exposed to *P. aeruginosa* (OD_600_=0.6) which kills *C. elegans* in a liquid medium. Twenty nematodes were used in each group. Representative survival curves of *C. elegans* following infection by *P. aeruginosa* in (A) liquid medium consisting of M9 buffer and *P. aeruginosa* culture, *Pseudomonas* phage, or heat-inactivated bacteria and (B, C, D) survival curves of *C. elegans* following infection with *P. aeruginosa* and treatment with phage Motto, therapeutic and prophylactic treatment. (B) Survival curves of *C. elegans* infected and treated with bacteria and phage ratio of 1:1, i.e., 10^5^ CFU/mL and 10^5^ PFU/mL. (C) Survival curves of *C. elegans* infected and treated with bacteria and phage ratio of 1:10, i.e., 10^5^ CFU/mL and 10^6^ PFU/mL. (D) Survival curves of *C. elegans* infected and treated with bacteria and phage ratio of 1:100, i.e., 10^5^ CFU/mL and 10^7^ PFU/mL. The survival curves were plotted using the Kaplan-Meier method, and the log-rank test was used to analyze the difference in survival rates in GraphPad Prism 9.0. A statistically significant difference (*p < 0*.*05*) was observed in the treatment groups.

In contrast, heat-inactivated bacteria did not cause any significant effect on the survival of nematodes, with 95% survival observed (Fig.7A). In the group in which the nematodes were exposed to phages but not to bacteria, no decrease in nematode survival was observed. To study the efficacy of the phage treatment, varying multiplicities of infection were tested, i.e., 1:1, 1:10, and 1:100. In the therapeutic treatment group, phage Motto was added after exposure of the nematode to the pathogen for 2 hours. Here, phage Motto was able to increase survival up to 90% within the 5 days observation time when treated at a 1:100 ratio (Fig.7D). The other ratios also increased the survival of nematodes, albeit not as significantly (Fig.7B, 7C). The more efficient activity was observed from the phage in the prophylactic treatment group, with nematode survival up to 80% after 5 days (Fig.7D). The phage treatment resulted in at least a 5-fold increase in nematode survival.

## Discussion

In this study, the *Pseudomonas* phage Motto was isolated from Cooum river water in Tamil Nadu, India. Genome analysis showed that the phage Motto is closely related to the *Drexlerviridae* family in the order *Caudoviricetes* (45). To date, *Pseudomonas* phages with a siphovirus-like morphology have been less well studied. Such phages have been found to have comparably narrow host ranges while also being predominantly lysogenic (46-49). However, our study reports a lytic phage with a broad host range.

Motto has a broad host range and is able to infect 64% (n = 32/50) of the tested clinical pseudomonal isolates. However, in most cases phage resistance is observed both, *in vitro* and *in vivo*, complicating treatment during therapy. Yet, phage-resistant mutants have been observed to often display decreased virulence and antibiotic resistance (50,51). Interestingly, we did not observe phage-resistant strains emerge under the conditions we tested, making phage Motto a promising candidate for therapy. We also observed rapid absorption and replication, as well as temperature and pH stability, which is considered advantageous for therapy.

We also addressed the question of the safety and efficacy of Motto in tissue culture and in an animal model. We observed no cytotoxic effect of Motto on HEK 293 and RAW 264.7 macrophages. In addition, phage Motto was able to increase the survival of *C. elegans* up to 90% at an MOI of 100 within the 5 days observation time using a recently established liquid-based test platform (42). At an MOI of 1, nematode survival was up to 80% at a lower MOI of only 1 (Fig.7). Overall, the phage treatment resulted in at least a 5-fold increase in nematode survival, illustrating the potential as a candidate for therapeutic applications.

Most lab-based studies of phages that are evaluated or considered for therapy, investigate planktonic cells and rarely deal with the issue of biofilms (52,53). However, *P. aeruginosa* often causes clinical complications due to the ability to form biofilms, often when present in chronic infections such as cystic fibrosis. Biofilms also can increase antibiotic resistance levels (54,55). Thus, phages that have the ability to disrupt biofilms are of particular interest, in order to use biofilm-degrading proteins such as depolymerases or the virus as a whole. Motto was able to reduce the biofilm significantly (Fig.4), similar to a previous report (53). The same study also reported the impact of exposure time, 3 h and 24 h, in reducing the biofilms. Importantly, our study showed the reduction of strong biofilms after 24 hours of treatment, in comparison to previous studies in which no anti-biofilm effect was observed for mucoid clinical strains which produced extensive biofilms (53,56).

We previously reported the complete genome of the phage Motto (57). Here, we provide details on the 84 ORFs that were predicted; 38 of which have assigned putative functions (S.table 2). The largest group is comprised of those that form the virus structure and morphogenesis proteins (such as chaperones, and tail measure proteins). We identified 11 proteins that are putatively interacting with, degrading or modifying nucleic acids of either the phage or the host, including a single-stranded DNA-binding protein (locus_tag=Motto_27), and putative DNA modifying proteins; these include a C-5 cytosine DNA methylase, a deoxynucleotide monophosphate kinase, a phosphoesterase, and a Dam methylase. Lysis-related proteins include a putative holin, a SAR endolysin, and a u-spanin. Interestingly, we also identified a putative super-infection exclusion protein (locus_tag=Motto_24).

The comparative analysis suggests that Motto represents a new species and genus within the family *Drexlerviridae*. To the best of our knowledge, this is the first T1-like phage characterized to infect *Pseudomonas* species.

## Conclusion

We isolated a new phage infecting *Pseudomonas aeruginosa*, Motto, which exhibits a broad host range, infecting more than 50% of the strains we tested (n = 50). In addition to the phage being physically stable and replicating rapidly, two properties make the bacteriophage a promising therapeutic candidate. Firstly, Motto activity against biofilms and secondly that no phage resistance was observed under the conditions we tested. These properties are highly advantageous for the treatment of *P. aeruginosa* infections employing phage therapy.

## Acknowledgement

The authors would also like to thank the VIT-TEM facility and the funding support provided by the VIT-SEED grant. The authors gratefully acknowledge Caenorhabditis Genetic Centre (CGC, University of Minnesota, MN, USA) which is funded by the NIH Office of Research Infrastructure Programs (P40 OD010440) for providing Bristol N2 (wild-type) used in this work.

## Author’s contribution

Conceptualization, P.M., and R.N.; methodology, P.M., M.M., D.T., and R.N.; validation, P.M., R.N., and R.T.; formal analysis, P.M., M.M., and B.L.; investigation, P.M., and R.N.; resources, N.E., R.T., R.N., D.T., and S.L.; writing-original draft preparation, P.M.; writing-review and editing, R.N., B.L., D.T., and S.L.; supervision, R.N., and S.L.; project administration, P.M., and S.L.; funding acquisition, P.M. All authors have read and agreed to the published version of the manuscript.

## Conference

A part of this research data was presented at the Second International Conference on Bacteriophage Research (ICBR), India in 2021.

